# Pathogenesis, transmission and response to re-exposure of SARS-CoV-2 in domestic cats

**DOI:** 10.1101/2020.05.28.120998

**Authors:** Angela M. Bosco-Lauth, Airn E. Hartwig, Stephanie M. Porter, Paul W. Gordy, Mary Nehring, Alex D. Byas, Sue VandeWoude, Izabela K. Ragan, Rachel M. Maison, Richard A. Bowen

**Affiliations:** College of Veterinary Medicine and Biomedical Sciences, Colorado State University, Fort Collins, CO

## Abstract

The pandemic caused by SARS-CoV-2 has reached nearly every country in the world with extraordinary person-to-person transmission. The most likely original source of the virus was spillover from an animal reservoir and subsequent adaptation to humans sometime during the winter of 2019 in Wuhan Province, China. Because of its genetic similarity to SARS-CoV-1, it is likely that this novel virus has a similar host range and receptor specificity. Due to concern for human-pet transmission, we investigated the susceptibility of domestic cats and dogs to infection and potential for infected cats to transmit to naïve cats. We report that cats are highly susceptible to subclinical infection, with a prolonged period of oral and nasal viral shedding that is not accompanied by clinical signs, and are capable of direct contact transmission to other cats. These studies confirm that cats are susceptible to productive SARS-CoV-2 infection, but are unlikely to develop clinical disease. Further, we document that cats develop a robust neutralizing antibody response that prevented re-infection to a second viral challenge. Conversely, we found that dogs do not shed virus following infection, but do mount an anti-viral neutralizing antibody response. There is currently no evidence that cats or dogs play a significant role in human exposure; however, reverse zoonosis is possible if infected owners expose their domestic pets during acute infection. Resistance to re-exposure holds promise that a vaccine strategy may protect cats, and by extension humans, to disease susceptibility.

## Introduction

The COVID-19 pandemic, caused by the SARS-CoV-2 (SARS2) coronavirus, originated in the Wuhan province of China, in late 2019 and within four months spread to nearly every country in the world. Sequence analysis and epidemiological investigations suggest that the virus was of animal-origin, possibly bat, and was first introduced into the human population via an intermediate animal host in the Huanan seafood market in Wuhan, China (Bogoch et al. 2020; Zhou et al. 2020). The virus quickly adapted to humans and human-to-human transmission became the almost immediate source of subsequent infections, with direct contact and aerosol droplets as the primary routes of infection (Li et al. 2020). Early indications suggested that SARS2, much like SARS-CoV-1 (SARS1), infects host cells by binding to the angiotensin-converting enzyme 2 (ACE2), a receptor that is expressed in many animal species, although notably not in mice or rats (Wan et al. 2020). Thus, while humans are almost certainly the sole source of infection to other humans, multiple early studies suggest other animals are susceptible to infection as well.

The first report of reverse zoonosis, or transmission from human to animal, was reported from Hong Kong, where a COVID patient’s dog tested PCR positive for SARS2 multiple times (Sit et al. 2020). In following weeks, other reports of domestic pets becoming infected following exposure to humans were documented, including another dog in Hong Kong and a cat with clinical disease in Belgium (Chini 2020). Serologic studies so far have failed to identify domestic dogs and cats as a primary source of human infection (Deng et al. 2020). Importantly, a survey of veterinary students with confirmed COVID infection was unable to identify antibodies in their pets (Temmen et al. 2020). Despite the low probability of pet-to-human or human-to-pet transmission, it remains important to clarify what role, if any, that domestic pets play in SARS2 transmission.

The first published study involving cat experimental infections showed that cats could become infected by SARS2 and could potentially transmit to other cats via aerosols, as defined by PCR positive fecal samples from cats in cages in the same room as directly infected cats. This study also described pathology and mortality in juvenile cats euthanized at 3 and 7 days post-infection (Shi et al. 2020) Additional communications described viral shedding and direct contact transmission in cats as well as seroconversion in cats exposed to infected humans (Halfmann et al. 2020; Zhang et al. 2020). The experiments described herein expand upon existing work by providing shedding kinetics in cats over time, assessing virus neutralization, seroconversion, and exploring transmission. Furthermore, to the author’s knowledge, this is the first report of protective immunity against SARS2 following repeated exposure. These studies indicate that cats may serve as a suitable animal model for studying SARS2 infection, and for furthering the development of vaccines and therapeutics for use in both animals and humans. We also confirm an earlier report that dogs do not replicate virus locally (Shi et al. 2020), but document evidence of anti-viral neutralizing activity in post-exposure canine sera. The role of cats in zoonotic transmission remains an open question, but relatively short duration of shedding and resistance to re-exposure suggests risk of this is very low, particularly when cats are kept indoors.

## Materials and Methods

### Virus

SARS2 virus strain WA1/2020WY96 was obtained from BEI Resources (Manassas, VA, USA), passaged twice in E6 Vero cells and stocks frozen at −80C in Dulbecco’s Modified Eagle Medium (DMEM) with 5% fetal bovine serum and antibiotics. Virus stock was titrated on Vero E6 cells using standard double overlay plaque assay (Kropinsky et al. 2008) and plaques counted 72 hours post-infection to determine plaque-forming units (pfu) per ml.

### Animals

Seven adult (1 male, 6 female, 5-8 year old) cats were obtained from a closed breeding colony held at Colorado State University in a pathogen-free environment in an Association for Assessment and Accreditation of Laboratory Animal Care (AAALAC) International accredited animal facility. Cats were screened negative for feline enteric coronavirus antibody prior to transfer. Three dogs (female, 5-6 years old) were obtained from Ridglan Farms (Blue Mounds, WI, USA). Cats and dogs were transferred to the Animal Disease Laboratory, an Animal Biosafety Level-3 (ABSL3) facility at Colorado State University, group housed and fed dry/wet food mix with access to water ad libitum. Animals were allowed several days to acclimate before temperature-sensing microchips (Lifechips, Destron-Fearing) were inserted subcutaneously in the dorsum. Baseline weights, body temperatures, clinical evaluation, and oral swabs were obtained prior to inoculation. All animals were in apparent good health at the onset of the study.

### Virus challenge

Cats were lightly anesthetized with 30-50 mg subcutaneous ketamine hydrochloride (Zetamine^™^) and dogs sedated with 1-3 mg xylazine. Virus diluted in phosphate buffered saline (PBS) was administered to both species via pipette into the nares (500ul/nare) for a total volume of 1ml; animals were observed until fully recovered from anesthesia. Virus back-titration was performed on E6 cells immediately following inoculation, confirming that cats received 3.0E5 pfu and dogs received 1.4E5 pfu.

### Sampling

#### Cat cohort 1 (n=3)

Oropharyngeal swabs were collected daily on days 1-5, 7, 10, and 14 post-infection using a polyester tipped swab applicator. Swabs were placed in BA-1 medium (Tris-buffered MEM containing 1% BSA) supplemented with gentamicin, amphotericin B and penicillin/streptomycin. Nasal flushes were performed on 1, 3, 5, 7, 10, and 14 days post-infection (DPI) by instilling 1ml BA-1 dropwise into the nares of awake or lightly anesthetized cats and collecting nasal discharge into a sterile petri dish by allowing the wash fluid to be sneezed out or dripped onto the dish. Blood (5 ml into serum separator tubes) was collected prior to inoculation and on days 7, 14, 21, 28, 35 and 42 post-infection. At 28 DPI, cats were re-challenged with 3e5 pfu of homologous virus. Oronasal sample collection was performed 1, 3, 5, 7, 10 and 14 after re-inoculation (days 29, 31, 33, 35, 38 and 42 post initial challenge), at which time cats were euthanized and tissues collected for histopathology.

#### Cat cohort 2 (n=4)

Two of the four cats were lightly anesthetized, and challenged with SARS2 as for Cohort 1. Forty-eight hours post-infection, two naïve cats were introduced into the room with the infected cats and sampled on the same schedule as before. The two directly challenge cats were euthanized on 5 DPI and the following tissues collected for virus isolation and histopathology: nasal turbinates, trachea, esophagus, mediastinal lymph node, lung, liver, spleen, kidney, small intestine, uterus, and olfactory bulb. Tissues were collected into BA-1 frozen at −80C and homogenized prior to plaque assay. Additional tissues collected for histopathology included heart, colon, pancreas, hemi-lung lobe, and mesenteric lymph nodes. Thoracic radiographs were also obtained for these two cats pre-challenge and just prior to euthanasia. The remaining two cats were euthanized at 30 DPI and necropsied; these cats will be referred to as contact cats hereafter.

#### Dogs (n=3)

Dogs were sampled at the same frequency, and using the same methods as cats in Cohort 1 for 42 days post-infection. Dogs were not re-challenged.

### Clinical observations

Body temperatures were recorded daily for the duration of the study using the thermal microchips. Cats and dogs were observed twice daily for the first seven days post-challenge and at least once daily for the duration of the study. Body weights were obtained weekly. Clinical evaluation included temperament, ocular discharge, nasal discharge, ptyalism, coughing/sneezing, dyspnea, diarrhea, lethargy, anorexia, moribund. None of the animals exhibited clinical signs of disease characterized by any of these symptoms at any time during the study.

### Viral assays

Virus isolation was performed on all oral swab, nasal flush and 5 DPI tissue samples by double overlay plaque assay on Vero E6 cells as previously described (Kropinsky et al. 2009). Briefly, 6-well plates with confluent monolayers of cells were washed once with PBS and inoculated with 100 ul of serial 10-fold dilutions of samples, incubated for 1 hour at 37°C, and overlaid with a 0.5% agarose in MEM containing 2% fetal bovine serum and antibiotics/antifungal agents. A second overlay with neutral red dye was added at 48 hours and plaques were counted at 72 hours. Viral titers were reported as the log10 plaque-forming units (pfu) per ml.

Plaque reduction neutralization assays (PRNT) were performed as previously described (Perera et al. 2020). Serum was heat-inactivated for 30 mins at 56°C, and two-fold dilutions prepared in BA-1 starting at a 1:5 dilution were aliquoted onto 96-well plates. An equal volume of virus was added to the serum dilutions and incubated for 1 hour at 37°C. Following incubation, serum-virus mixtures were plated onto Vero E6 plates as described for virus isolation assays. Antibody titers were recorded as the reciprocal of the highest dilution in which >90% of virus was neutralized.

### ELISA

Serum samples from cats were heat inactivated and tested by plaque assay to verify samples were noninfectious prior to conducting ELISA analysis. Positive control antibodies to the receptor-binding domain (RBD) and full-length spike protein were human MAb CR3022 antibody (Absolute Antibody, Oxford UK) and human IgG whole molecule (Jackson Immuno Research, West Grove PA, USA). Positive control for the nucleocapsid ELISA was SARS-CoV nucleoprotein rabbit monoclonal antibody (Sino Biological Inc, Beijing, China). Negative controls were reagent grade human sera (compared to Mab CR3022). Cat serum from specific pathogen free, naïve experimental animals, and field isolate bioarchived samples obtained prior to 2019 (Carver et al, 2005, Sprague et al, 2018). ELISA protocols were adapted from protocols for SARS CoV-2 ELISA described by Amanat et al., 2020. ELISA plates (Thermo) were coated at 2ug/ml with spike glycoprotein Receptor Binding Domain (RBD) from SARS-CoV-2, WuHan-Hu-1 recombinant from HEK293T cells (BEI), or Spike glycoprotein (Stabilized) from SARS-CoV-2, Wuhan-Hu-1, recombinant from Baculovirus (BEI). SARS CoV-2 nucleocapsid protein was a gift of Dr. Brian Geiss. Prior to running experimental cat sera, the assay was optimized using positive and negative control sera described above (data not shown.) Samples and controls were diluted 1:50 in ELISA diluent (1X PBS, tween, milk powder) and run in duplicate. Human sera controls were developed using anti-human IgG HRP (Thermo), cat sera was developed using anti-cat IgG HRP (Thermo) or anti-cat IgM (Novus Biologicals) and rabbit mAb SARS-CoV NP was detected by anti-rabbit IgG HRP (Thermo). Secondary antibodies were diluted 1:3000 and-SigmaFast OPD was prepared in WFI and added to wells. Plates were read at 490nm using a Multiskan® Spectrum spectrophotometer (Thermo Fisher). The mean of negative control sera OD490 plus three times the standard deviation of the negative control readings were used to determine cut off values for each plate.

### qRT-PCR

Plaques were picked from culture plates from each cat to confirm SARS2 viral shedding. RNA extractions were performed per the manufacturer’s instructions using Qiagen QiaAmp Viral RNA mini kits. RT-PCR was performed as recommended using the E_Sarbeco primer probe sequence as described by Corman and colleagues (2020) and the Superscript III Platinum One-Step qRT-PCR system (Invitrogen), with the following modification; the initial reverse transcription was at 50°C. Standard curves were obtained by serial dilution of stock viral RNA from the original WA1/2020WY96 SARS2 isolate.

### Histopathology

Tissues from cats were fixed in 10% buffered formalin for 12 days and transferred to 70% ethanol prior to sectioning for H&E staining. Slides were read by a board certified veterinary pathologist.

## Results

### Clinical Disease

None of the cats in either cohort displayed any clinical signs of disease and remained afebrile throughout the study. Body weights were maintained over time. Radiographs were taken pre-challenge and at 5 DPI just prior to euthanasia for the experimentally inoculated cats in Cohort 2. No evidence of lung involvement or any other radiographically-detectable abnormalities were noted (images not shown). Similarly, dogs inoculated with SARS2 remained clinically normal and afebrile.

### Viral shedding

In Cohort 1, all three cats shed virus both orally and nasally for up to 5 days post-infection, with peak titers achieved from nasal shedding at day 3. Nasal titers were approximately 1 log higher than oral swabs collected at the same time (Figure 1). There was some variability in titer over the course of infection that is likely attributable to sample collection (i.e. quality of sneezes), but overall the data demonstrates clear presence of infectious virus in both the nasal cavity and the oropharynx for multiple days post-infection. In Cohort 2, the inoculated cats shed virus for 5 days post-infection both orally and nasally, with a similar pattern to Cohort 1. The contact cats, however, shed infectious virus orally by 24 hours post-exposure and the duration of shedding was prolonged compared to the inoculated cats, with peak shedding occurring at 7 days post-exposure (Figure 1). Virus was isolated from trachea, nasal turbinates, and esophagus from cats in Cohort 2 necropsied on day 5. Infectious virus was not found in the lung or other organs of either cat. Viral shedding was not detected from any of the dogs at any point post-infection.

**Figure 1:**
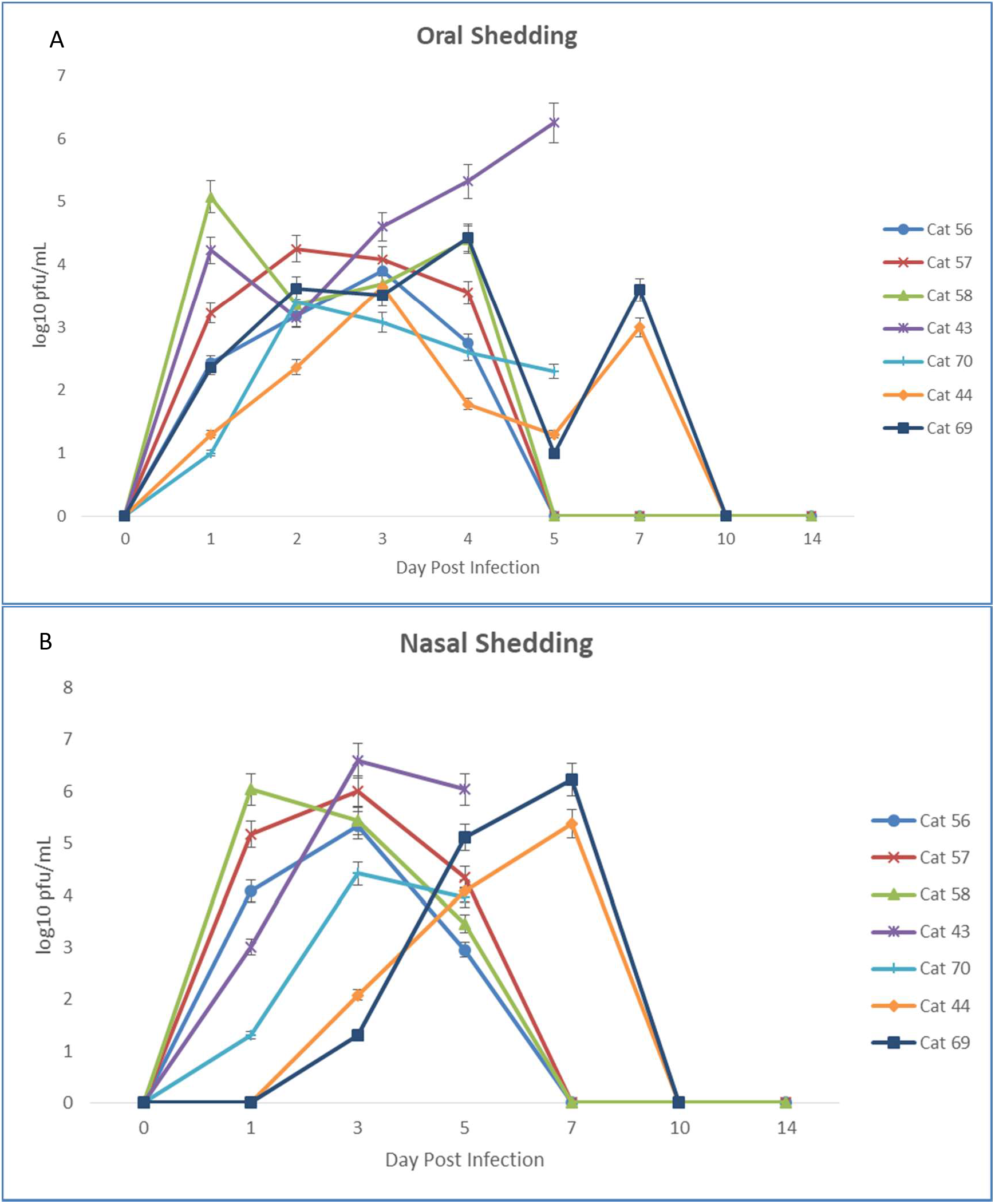
Inoculation and exposure with SARS2 leads to oral and nasal shedding in cats. SARS2 virus is detected by plaque assay from (A) oral and (B) nasal secretions of cats 1-5 days post infection. Viral titers expressed as log10pfu/ml. Cats 56, 57, and 58 represent Cohort 1. Cats 43, 70, 44, and 69 represent Cohort 2. Cats 43 and 70 were euthanized on 5 DPI. Cats 44 and 69 were introduced to the infected cats in Cohort 2 on 2 DPI.

### Pathology

Gross lesions were not observed in any of the necropsied cats. Histologically, in both cats sacrificed at 5 DPI from Cohort 1, moderate ulcerative, suppurative lymphoplasmacytic rhinitis was observed in the nasal turbinates along with mild lymphoplasmacytic tracheitis. These cats also had minimal alveolar histiocytosis with edema and one of the cats had rare type II pneumocyte hyperplasia (Figure 2). All three cats from Cohort 1 sacrificed 42 DPI had mild lung changes, including mild interstitial lymphocytic pneumonia with peribronchiolar and perivascular lymphocytic cuffing and alveolar histiocytosis. Two of these cats also had minimal tracheitis, but largely the lesions in the upper respiratory tract appear decreased in comparison to the early timepoint cats, while lung pathology was more evident in these animals compared to those sacrificed during acute infection. Dogs were not euthanized at the time of this report.

**Figure 2:**
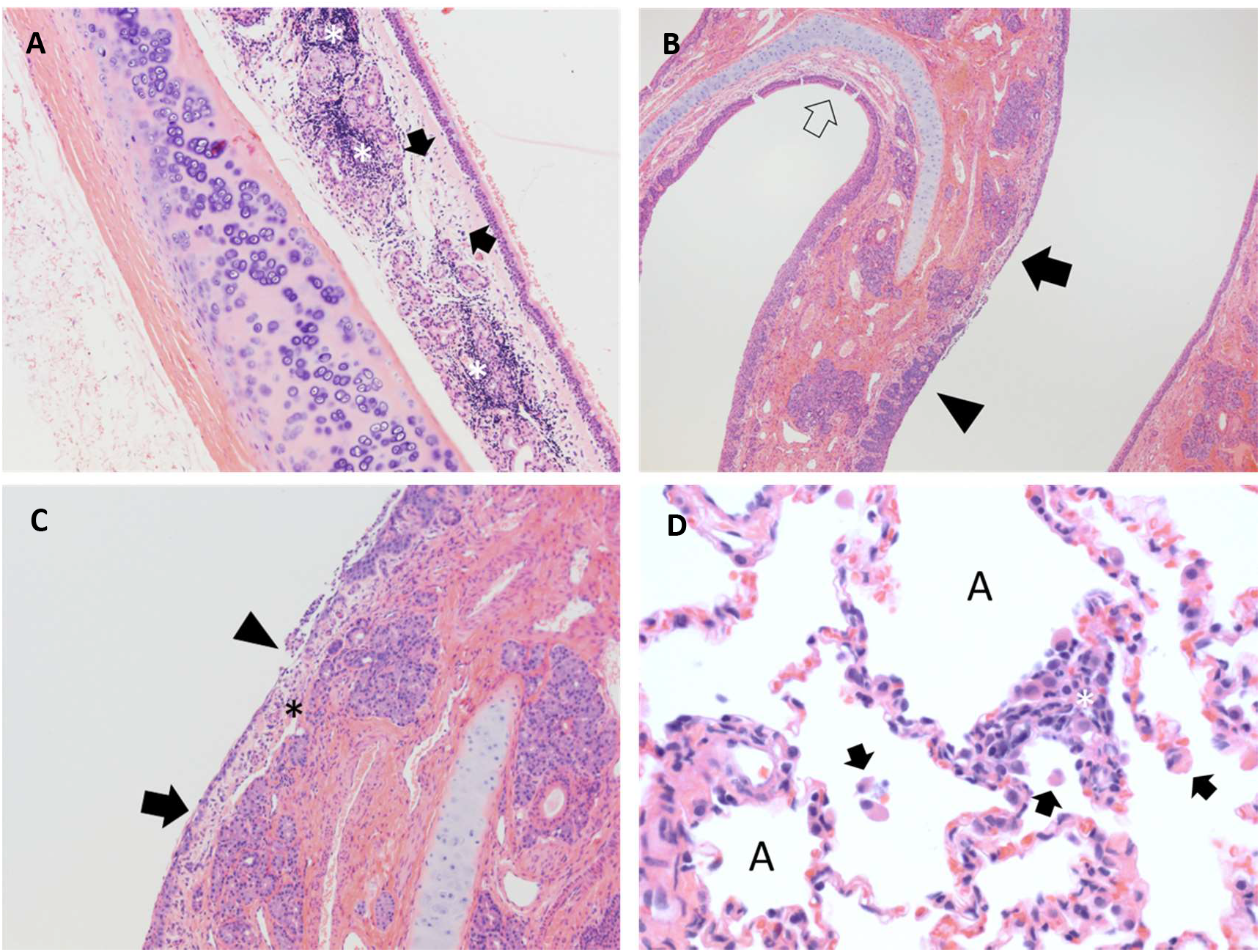
SARS2 exposure results in acute upper respiratory inflammation and mild lung infiltrates during later courses of infection. Panel A: **Cat 43, trachea 5 DPI.** The submucosa is expanded by edema (arrow) and abundant lymphocytic inflammatory infiltrates (asterisks) which dissect and disrupt submucosal glands. Hematoxylin & eosin stain. 100x magnification. Panel B: **Cat 70, nasal turbinates 5 DPI.** Normal thickness respiratory mucosa is present in the section (open arrow). Nasal respiratory epithelium ranges from hyperplastic (filled black arrow) to ulcerated (arrowhead). The submucosa in regions of ulceration is edematous and infiltrated by scattered neutrophils and mononuclear cells. Hematoxylin & eosin stain. 40x magnification. Panel C: **Cat 70, nasal turbinates, 5 DPI.** Nasal respiratory epithelium ranges from attenuated (arrow) to ulcerated (arrowhead) with overlying remnant cellular debris. The submucosa (asterisk) in regions of ulceration is edematous and infiltrated by scattered neutrophils and mononuclear cells. Hematoxylin & eosin stain. 100x magnification. Panel D: **Cat 56, lung, 42 DPI.** Alveolar spaces (A) contain scattered mononuclear cells (arrow). The alveolar wall is expanded by mixtures of mononuclear cells and occasional neutrophils (asterisk). Hematoxylin & eosin stain. 400x magnification.

### Seroconversion

Cats in both Cohort 1 and the direct contact cats developed neutralizing activity as measured by PRNT as early as 7 DPI. Neutralizing titers in all cats reached or exceeded 1:2560 by 14 DPI and either maintained or increased in titer between 28 and 42 DPI. Cats re-challenged at 28 DPI displayed a moderate increase in PRNT titer in the 14 days following exposure (Table 1). Dogs developed neutralizing antibodies by 14 DPI and peaked at 21 DPI with titers between 1:40-1:80 (Table 1).

**Table 1:** PRNT90 values for cats and dogs.

| Animal | 0 DPI | 7 DPI | 14 DPI | 21 DPI | 28 DPI | 42 DPI |
| --- | --- | --- | --- | --- | --- | --- |
| Cat 56 (Cohort 1) | <50% | 320 | 5120 | 2560 | 2560 | 10240 |
| Cat 57 (Cohort 1) | <50% | 80 | 5120 | 2560 | 2560 | 5120 |
| Cat 58 (Cohort 1) | <50% | 80 | 2560 | 2560 | 1280 | 5120 |
| Cat 43 (Cohort 2) | <50% |  |  |  |  |  |
| Cat 70 (Cohort 2) | <50% |  |  |  |  |  |
| Cat 44 (Cohort 2) | <50% | NT | 2560 | 5120 | 5120 |  |
| Cat 69 (Cohort 2) | <50% | NT | 2560 | 1280 | 10240 |  |
| Dog 46 | <50% | <10 | 10 | 40 | 40 | 80 |
| Dog 47 | <50% | 10 | 80 | 20 | 20 | 40 |
| Dog 48 | <50% | <10 | 20 | 40 | 20 | 20 |

IgG antibodies responses exceeding OD490 cut off values were detected at day 7 PI against both the complete spike glycoprotein and RBD in all experimentally inoculated cats, and seroconversion against NP was detected in 2 of 3 cats at this time. By day 14 all five cats had OD values that neared upper limit of detection in the Spike ELISA; RBD and NP OD saturation was obtained by day 21 and did not increase following re-exposure (Figure 3). Rates of seroconversion and absorbance levels were similar between contact cats and experimentally infected cats. Seroconversion to spike protein was most rapid and robust, and the specificity of response to RBD exceeded that of NP. Seroconverted cat OD values for all three antigens exceeded absorbances of SPF or field domestic cats, and background was highest for NP. IgM antibodies against RBD were detected at days 7 and 14 but not at day 28. IgG responses were much more robust than IgM (Figure 3). ELISA assays were not performed on dog sera.

**Figure 3.**
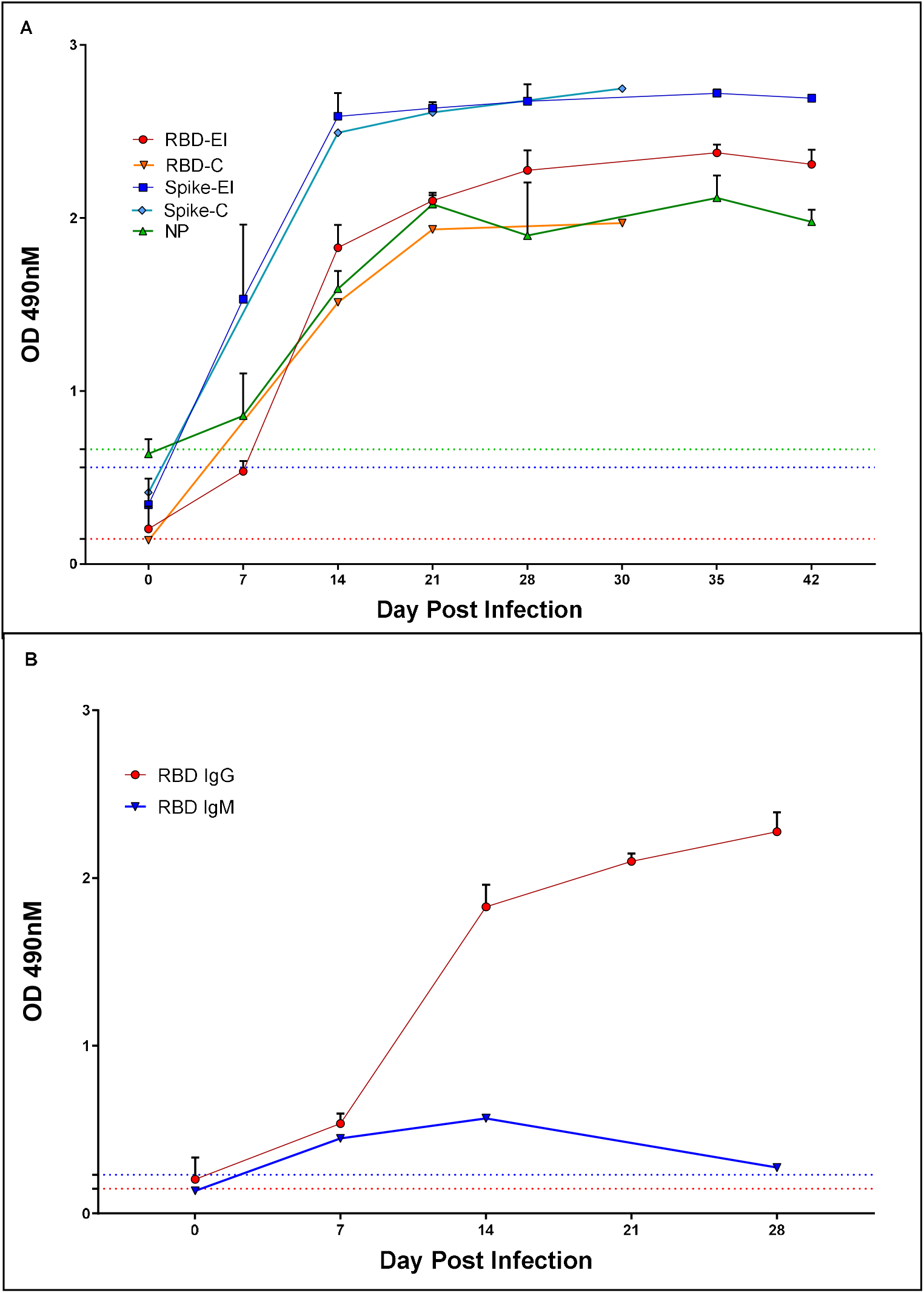
Cats infected with SARS CoV-1 rapidly develop antibodies against viral antigens. (A) Sera from cats with intranasal inoculation of SARS CoV-2 (n=3, ‘EI’) or exposed to inoculated cats (n=2, ‘C’) were evaluated for seroreactivity to receptor binding domain (RBD), Spike, or nucleocapsid protein (NP) for 30-42 days post exposure. IgG Reactivity to Spike and RBD was evident at day 7, and all animals had clearly seroconverted by day 14. (B) IgM against RBD was transiently detected at low levels relative to IgG on days 7 and 14 post exposure (experimentally inoculated animals, n=3). Bars represent 1 SE of the mean.

### Reinfection

Re-challenged cats in Cohort 1 were sampled for oral and nasal shedding for 7 days post-exposure by viral isolation, and shedding was not detected in any cat at any timepoint following rechallenge.

## Discussion

The COVID19 pandemic represents the first global pandemic of an emerging zoonotic disease in this century. The SARS2 virus is one of three emergent zoonotic coronaviruses capable of causing significant disease in humans in the last two decades, following SARS1 and MERS (Guarner 2020). The overall trend of disease emergence favors viral spillover from animals to human, and land use and wildlife encroachment are just two of the factors contributing to this phenomenon (Olival et al. 2017). The continued presence of live animal markets provides optimal conditions for emergence of zoonoses (Wang and Eaton, 2007). As with SARS1 and MERS, SARS2 is of probable bat origin based on phylogenetic analysis (Zhou et al. 2020), but unlike its predecessors, SARS2 has rapidly evolved for highly efficient human-to-human transmission (Chan et al et al. 2020). While animals, including domestic animals and pets, are frequently implicated as the source of emerging pathogens, reverse zoonosis of SARS2 is more probable, as human cases are far more prevalent than domestic animals. Similar results were seen with SARS1, where domestic cats exposed to the virus by infected humans became infected, and cats experimentally infected shed virus for several days (Martina et al. 2003, van den Brand et al. 2008) There have been several cases of pets becoming infected by SARS2 following exposure to infected humans in New York, Hong Kong, Belgium, Germany, Spain, France and Russia (Sit et al. 2020; as communicated by ProMed). Other animal exposures from infected humans include farmed mink, which appear to display respiratory symptoms following infection (ProMed). In several of these cases, including nondomestic felids at the Bronx Zoo and pet cats in New York and Europe, animals displayed signs of respiratory disease and/or conjunctivitis. None of the cats or dogs in this study exhibited any clinical signs of disease, but individual animal health status, age and comorbidities may be responsible for this variability. Pathological changes in cats suggest that mild subclinical disease in otherwise healthy animals occurs but is not recognizable symptomatically. This is not altogether different from human infections, where the majority of cases are relatively mild but more severe disease tends to occur in older patients with significant comorbidities (Nikolich-Zugich et al. 2020). In a recent serosurvey of cats in Wuhan, China, nearly 14.7% of sampled animals were seropositive for SARS2 by RBD ELISA, suggesting that the cat population in areas with high human transmission is also likely to be exposed to the virus (Zhang et al. 2020) Considering that the number of human infections have reached the millions and yet only a handful of animals have tested PCR positive, it seems unlikely that domestic pets are a significant source of infection or are at serious risk for developing severe disease. Importantly, infected cats shed for no more than 5 days following exposure, suggesting that cats, if exposed to infected humans, will develop and clear infection rapidly. In comparison, humans typically have an incubation period of approximately 5 days and can shed virus for more than three weeks (Lauer et al. 2020, Noh et al. 2020). Thus, if symptomatic humans follow appropriate quarantine procedures and stay home with their pets, there is minimal risk of a potentially exposed cat infecting another human. Infected pet cats should not be allowed outdoors to prevent potential risk of spreading infection to other outdoor cats. More research into the susceptibility of wildlife species and potential for establishment of infection in outdoor cat populations is necessary to identify risk factors and mitigation strategies to prevent establishment of reservoir infections in feral cats.

The development of animal models for studying SARS-2 is an important step in research methodologies. Rhesus macaques, hamsters, and ferrets are all suitable models for replicating asymptomatic or mildly clinical disease, and, while not often used as a traditional animal model, this work demonstrates that cats may serve as an alternative model (Kim et al. 2020, Chan et al. 2020, Munster et al. 2020) The cats in this study developed subclinical pathological changes in the upper respiratory tract early in the course of infection with more lower respiratory tract pathology later following viral clearance, which suggests that, while subclinical, viral infection of cats is not completely benign, and may make their utility as an animal model more relevant to mild human disease. Additionally, the relatively high-titer viral shedding produced by cats and the rapidity of transmission may make them an ideal model for simulation of aerosols. As such, cat models may be quite useful for understanding the shed/spread kinetics of SARS2. Perhaps most importantly, cats develop significant neutralizing antibody titers, and are resistant to re-infection, which could prove a useful measurement for subsequent vaccine trials for both human and animal vaccine candidates.

## Acknowledgements

The authors thank Todd Bass and the histology lab at Colorado State University for preparation of tissue cassettes and slides for histopathology and Dr. Brian Geiss for providing the SARS-2 nucleocapsid protein.

## Author contributions

**Angela Bosco-Lauth:** Conceptualization, Methodology, Data curation, Writing. **Airn Hartwig:** Conceptualization, Methodology, Data curation, Writing. **Stephanie Porter:** Conceptualization, Methodology, Data curation, Writing. **Paul Gordy:** Investigation, Review & Editing. **Mary Nehring:** Investigation, Review & Editing. **Alex Byas:** Formal analysis, Review & Editing. **Sue VandeWoude:** Resources, Review & Editing. **Izabela Ragan:** Review & Editing. **Rachel Maison:** Review & Editing. **Richard Bowen:** Project administration; Review & Editing.

## Data and materials availability

SARS-Related Coronavirus 2, Isolate USA-WA1/2020 (NR-52281) was deposited by the Centers for Disease Control and Prevention and obtained through BEI Resources, NIAID, NIH. The following reagents were produced under HHSN272201400008C and obtained through BEI Resources, NIAID, NIH: Spike Glycoprotein Receptor Binding Domain (RBD) from SARS-Related Coronavirus 2, Wuhan-Hu-1, Recombinant from HEK293 Cells, NR-52306, and Spike Glycoprotein (Stabilized) from SARS-Related Coronavirus 2, Wuhan-Hu-1, Recombinant from Baculovirus, NR-52396. All data is available in the main text or the supplementary materials.

## References

1. I. Bogoch, A. Watts, A. Thomas-Bachli, C. Huber, M. U. G. Kraemer, K. Khan, Pneumonia of unknown aetiology in Wuhan, China: potential for international spread via commercial air travel. Journal of Travel Medicine. 27, taaa008 (2020).

2. P. Zhou, X.-L. Yang, X.-G. Wang, B. Hu, L. Zhang, W. Zhang, H.-R. Si, Y. Zhu, B. Li, C.-L. Huang, H.-D. Chen, J. Chen, Y. Luo, H. Guo, R.-D. Jiang, M.-Q. Liu, Y. Chen, X.-R. Shen, X. Wang, X.-S. Zheng, K. Zhao, Q.-J. Chen, F. Deng, L.-L. Liu, B. Yan, F.-X. Zhan, Y.-Y. Wang, G.-F. Xiao, Z.-L. Shi, A pneumonia outbreak associated with a new coronavirus of probable bat origin. Nature (2020), doi:10.1038/s41586-020-2012-7.

3. Q. Li, X. Guan, P. Wu, X. Wang, L. Zhou, Y. Tong, R. Ren, K. S. M. Leung, E. H. Y. Lau, J. Y. Wong, X. Xing, N. Xiang, Y. Wu, C. Li, Q. Chen, D. Li, T. Liu, J. Zhao, M. Liu, W. Tu, C. Chen, L. Jin, R. Yang, Q. Wang, S. Zhou, R. Wang, H. Liu, Y. Luo, Y. Liu, G. Shao, H. Li, Z. Tao, Y. Yang, Z. Deng, B. Liu, Z. Ma, Y. Zhang, G. Shi, T. T. Y. Lam, J. T. Wu, G. F. Gao, B. J. Cowling, B. Yang, G. M. Leung, Z. Feng, Early Transmission Dynamics in Wuhan, China, of Novel Coronavirus–Infected Pneumonia. N Engl J Med. 382, 1199–1207 (2020).

4. Y. Wan, J. Shang, R. Graham, R. S. Baric, F. Li, Receptor Recognition by the Novel Coronavirus from Wuhan: an Analysis Based on Decade-Long Structural Studies of SARS Coronavirus. J Virol. 94, e00127–20, /jvi/94/7/JVI.00127-20.atom (2020).

5. T. H. C. Sit, C. J. Brackman, S. M. Ip, K. W. S. Tam, P. Y. T. Law, E. M. W. To, V. Y. T. Yu, L. D. Sims, D. N. C. Tsang, D. K. W. Chu, R. A. P. M. Perera, L. L. M. Poon, M. Peiris, Infection of dogs with SARS-CoV-2. Nature (2020), doi:10.1038/s41586-020-2334-5.

6. M. Chini. Coronavirus: Belgian woman infected her cat [Internet]. The Brussels Times. 2020 [cited 2020 Apr 1]. https://www.brusselstimes.com/all-news/belgium-all-news/103003/coronavirus-belgian-woman-infected-her-cat/.

7. J. Deng, Y. Jin, Y. Liu, J. Sun, L. Hao, J. Bai, T. Huang, D. Lin, Y. Jin, K. Tian, Serological survey of SARS-CoV-2 for experimental, domestic, companion and wild animals excludes intermediate hosts of 35 different species of animals. Transbound Emerg Dis (2020) in press, doi:10.1111/tbed.13577

8. S. Temmam, A. Barbarino, D. Maso, S. Behillil, V. Enouf, C. Huon, A. Jaraud, L. Chevallier, M. Backovic, P. Pérot, P. Verwaerde, L. Tiret, S. van der Werf, M. Eloit, “Absence of SARS-CoV-2 infection in cats and dogs in close contact with a cluster of COVID-19 patients in a veterinary campus” (preprint, Microbiology, 2020), doi:10.1101/2020.04.07.029090.

9. J. Shi, Z. Wen, G. Zhong, H. Yang, C. Wang, B. Huang, R. Liu, X. He, L. Shuai, Z. Sun, Y. Zhao, P. Liu, L. Liang, P. Cui, J. Wang, X. Zhang, Y. Guan, W. Tan, G. Wu, H. Chen, Z. Bu, Susceptibility of ferrets, cats, dogs, and other domesticated animals to SARS–coronavirus 2. Science, eabb7015 (2020).

10. P. J. Halfmann, M. Hatta, S. Chiba, T. Maemura, S. Fan, M. Takeda, N. Kinoshita, S. Hattori, Y. Sakai-Tagawa, K. Iwatsuki-Horimoto, M. Imai, Y. Kawaoka, Transmission of SARS-CoV-2 in Domestic Cats. N Engl J Med, NEJMc2013400 (2020).

11. Q. Zhang, H. Zhang, K. Huang, Y. Yang, X. Hui, J. Gao, X. He, C. Li, W. Gong, Y. Zhang, C. Peng, X. Gao, H. Chen, Z. Zou, Z. Shi, M. Jin, “SARS-CoV-2 neutralizing serum antibodies in cats: a serological investigation” (preprint, Microbiology, 2020), doi:10.1101/2020.04.01.021196.

12. J. Guarner, Three Emerging Coronaviruses in Two Decades. American Journal of Clinical Pathology. 153, 420–421 (2020).

13. K. J. Olival, P. R. Hosseini, C. Zambrana-Torrelio, N. Ross, T. L. Bogich, P. Daszak, Host and viral traits predict zoonotic spillover from mammals. Nature. 546, 646–650 (2017).

14. L.-F. Wang, B. T. Eaton, in Wildlife and Emerging Zoonotic Diseases: The Biology, Circumstances and Consequences of Cross-Species Transmission, J. E. Childs, J. S. Mackenzie, J. A. Richt, Eds. (Springer Berlin Heidelberg, Berlin, Heidelberg, 2007; http://link.springer.com/10.1007/978-3-540-70962-61_3), vol. 315 of Current Topics in Microbiology and Immunology, pp. 325–344.

15. J. F.-W. Chan, S. Yuan, K.-H. Kok, K. K.-W. To, H. Chu, J. Yang, F. Xing, J. Liu, C. C.-Y. Yip, R. W.-S. Poon, H.-W. Tsoi, S. K.-F. Lo, K.-H. Chan, V. K.-M. Poon, W.-M. Chan, J. D. Ip, J.-P. Cai, V. C.-C. Cheng, H. Chen, C. K.-M. Hui, K.-Y. Yuen, A familial cluster of pneumonia associated with the 2019 novel coronavirus indicating person-to-person transmission: a study of a family cluster. The Lancet. 395, 514–523 (2020).

16. J. M. A. van den Brand, B. L. Haagmans, L. Leijten, D. van Riel, B. E. E. Martina, A. D. M. E. Osterhaus, T. Kuiken, Pathology of Experimental SARS Coronavirus Infection in Cats and Ferrets. Vet Pathol. 45, 551–562 (2008).

17. B. E. E. Martina, B. L. Haagmans, T. Kuiken, R. A. M. Fouchier, G. F. Rimmelzwaan, G. van Amerongen, J. S. M. Peiris, W. Lim, A. D. M. E. Osterhaus, SARS virus infection of cats and ferrets. Nature. 425, 915–915 (2003).

18. J. Nikolich-Zugich, K. S. Knox, C. T. Rios, B. Natt, D. Bhattacharya, M. J. Fain, SARS-CoV-2 and COVID-19 in older adults: what we may expect regarding pathogenesis, immune responses, and outcomes. GeroScience. 42, 505–514 (2020).

19. S. A. Lauer, K. H. Grantz, Q. Bi, F. K. Jones, Q. Zheng, H. R. Meredith, A. S. Azman, N. G. Reich, J. Lessler, The Incubation Period of Coronavirus Disease 2019 (COVID-19) From Publicly Reported Confirmed Cases: Estimation and Application. Annals of Internal Medicine. 172, 577–582 (2020).

20. J. Y. Noh, J. G. Yoon, H. Seong, W. S. Choi, J. W. Sohn, H. J. Cheong, W. J. Kim, J. Y. Song, Asymptomatic infection and atypical manifestations of COVID-19: comparison of viral shedding duration. Journal of Infection, S0163445320303108 (2020).

21. Y.-I. Kim, S.-G. Kim, S.-M. Kim, E.-H. Kim, S.-J. Park, K.-M. Yu, J.-H. Chang, E. J. Kim, S. Lee, M. A. B. Casel, J. Um, M.-S. Song, H. W. Jeong, V. D. Lai, Y. Kim, B. S. Chin, J.-S. Park, K.-H. Chung, S.-S. Foo, H. Poo, I.-P. Mo, O.-J. Lee, R. J. Webby, J. U. Jung, Y. K. Choi, Infection and Rapid Transmission of SARS-CoV-2 in Ferrets. Cell Host & Microbe, S1931312820301876 (2020).

22. V. J. Munster, F. Feldmann, B. N. Williamson, N. van Doremalen, L. Pérez-Pérez, J. Schulz, K. Meade-White, A. Okumura, J. Callison, B. Brumbaugh, V. A. Avanzato, R. Rosenke, P. W. Hanley, G. Saturday, D. Scott, E. R. Fischer, E. de Wit, Respiratory disease in rhesus macaques inoculated with SARS-CoV-2. Nature (2020), doi:10.1038/s41586-020-2324-7.

23. J. F.-W. Chan, A. J. Zhang, S. Yuan, V. K.-M. Poon, C. C.-S. Chan, A. C.-Y. Lee, W.-M. Chan, Z. Fan, H.-W. Tsoi, L. Wen, R. Liang, J. Cao, Y. Chen, K. Tang, C. Luo, J.-P. Cai, K.-H. Kok, H. Chu, K.-H. Chan, S. Sridhar, Z. Chen, H. Chen, K. K.-W. To, K.-Y. Yuen, Simulation of the clinical and pathological manifestations of Coronavirus Disease 2019 (COVID-19) in golden Syrian hamster model: implications for disease pathogenesis and transmissibility. Clinical Infectious Diseases, ciaa325 (2020).

24. A. M. Kropinski, A. Mazzocco, T. E. Waddell, E. Lingohr, R. P. Johnson, in Bacteriophages, M. R. J. Clokie, A. M. Kropinski, Eds. (Humana Press, Totowa, NJ, 2009; http://link.springer.com/10.1007/978-1-60327-164-6_7), vol. 501 of Methods in Molecular Biology, pp. 69–76.

25. R. A. Perera, C. K. Mok, O. T. Tsang, H. Lv, R. L. Ko, N. C. Wu, M. Yuan, W. S. Leung, J. M. Chan, T. S. Chik, C. Y. Choi, K. Leung, K. H. Chan, K. C. Chan, K.-C. Li, J. T. Wu, I. A. Wilson, A. S. Monto, L. L. Poon, M. Peiris, Serological assays for severe acute respiratory syndrome coronavirus 2 (SARS-CoV-2), March 2020. Eurosurveillance. 25 (2020), doi:10.2807/1560-7917.ES.2020.25.16.2000421.

26. S. Carver, S. N. Bevins, M. R. Lappin, E. E. Boydston, L. M. Lyren, M. Alldredge, K. A. Logan, L. L. Sweanor, S. P. D. Riley, L. E. K. Serieys, R. N. Fisher, T. W. Vickers, W. Boyce, R. McBride, M. C. Cunningham, M. Jennings, J. Lewis, T. Lunn, K. R. Crooks, S. VandeWoude, Pathogen exposure varies widely among sympatric populations of wild and domestic felids across the United States. Ecol Appl. 26, 367–381 (2016).

27. W. Sprague, R. Troyer, X. Zheng, B. Wood, M. Macmillan, S. Carver, S. VandeWoude, Prior Puma Lentivirus Infection Modifies Early Immune Responses and Attenuates Feline Immunodeficiency Virus Infection in Cats. Viruses. 10, 210 (2018).

28. V. M. Corman, O. Landt, M. Kaiser, R. Molenkamp, A. Meijer, D. K. Chu, T. Bleicker, S. Brünink, J. Schneider, M. L. Schmidt, D. G. Mulders, B. L. Haagmans, B. van der Veer, S. van den Brink, L. Wijsman, G. Goderski, J.-L. Romette, J. Ellis, M. Zambon, M. Peiris, H. Goossens, C. Reusken, M. P. Koopmans, C. Drosten, Detection of 2019 novel coronavirus (2019-nCoV) by real-time RT-PCR. Eurosurveillance. 25 (2020), doi:10.2807/1560-7917.ES.2020.25.3.2000045.

